# The immune cell landscape in kidneys of lupus nephritis patients

**DOI:** 10.1101/363051

**Authors:** Arnon Arazi, Deepak A. Rao, Celine C. Berthier, Anne Davidson, Yanyan Liu, Paul J. Hoover, Adam Chicoine, Thomas M. Eisenhaure, A. Helena Jonsson, Shuqiang Li, David J. Lieb, Edward P. Browne, Akiko Noma, Danielle Sutherby, Scott Steelman, Dawn E. Smilek, Patti Tosta, William Apruzzese, Elena Massarotti, Maria Dall’Era, Meyeon Park, Diane L. Kamen, Richard A. Furie, Fernanda Payan-Schober, Jill P. Buyon, Michelle A. Petri, Chaim Putterman, Kenneth C. Kalunian, E. Steve Woodle, James A. Lederer, David A. Hildeman, Chad Nusbaum, David Wofsy, Matthias Kretzler, Jennifer H. Anolik, Michael B. Brenner, the Accelerating Medicines Partnership in RA/SLE network, Nir Hacohen, Betty Diamond

**Affiliations:** Broad Institute of MIT and Harvard, Cambridge MA 02142; Division of Rheumatology, Allergy, Immunology, Department of Medicine, Brigham and Women’s Hospital, Harvard Medical School, Boston MA 02115; Internal Medicine, Department of Nephrology, University of Michigan, Ann Arbor, MI, USA 48109; Center for Autoimmune and Musculoskeletal Diseases, The Feinstein Institute for Medical Research, Northwell Health, Manhasset, NY 11030; Lupus Nephritis Trials Network, University of California San Francisco, San Francisco, CA 94107; Immune Tolerance Network, University of California San Francisco, San Francisco, CA 94107; Rheumatology Division, University of California San Francisco, San Francisco, CA 94107; Division of Nephrology, University of California, San Francisco, San Francisco, CA 94107; Division of Rheumatology and Immunology, Medical University of South Carolina, Charleston, SC 29425; Division of Rheumatology, Northwell Health, Great Neck, NY 11021; University of North Carolina Kidney Center, Division of Nephrology and Hypertension, Department of Medicine, UNC School of Medicine, Chapel Hill, NC, 27599; Department of Medicine, Division of Rheumatology, New York University School of Medicine, New York, NY, 10016; Division of Rheumatology, Johns Hopkins University, Baltimore, MD, 21287; Division of Rheumatology and Department of Microbiology and Immunology, Albert Einstein College of Medicine and Montefiore Medical Center, Bronx, NY, 10461; University of California San Diego School of Medicine, La Jolla, California, 92093; Division of Transplantation, Department of Surgery, University of Cincinnati College of Medicine, Cincinnati, OH 45257; Department of Surgery, Brigham and Women’s Hospital, Harvard Medical School, Boston, MA 02115; Department of Pediatrics, University of Cincinnati, Cincinnati, OH 45257; Division of Immunobiology, Cincinnati Children’s Hospital Medical Center, Cincinnati, OH 45229; Department of Medicine, Division of Allergy, Immunology, and Rheumatology, University of Rochester Medical Center, Rochester, NY 14642

**Author notes:** These authors contributed equally: Arnon Arazi, Deepak A. Rao and Celine C. Berthier.

## Abstract

Lupus nephritis is a potentially fatal autoimmune disease, whose current treatment is ineffective and often toxic. To gain insights into disease mechanisms, we analyzed kidney samples from lupus nephritis patients and healthy controls using single-cell RNA-seq. Our analysis revealed 21 subsets of leukocytes active in disease, including multiple populations of myeloid, T, NK and B cells, demonstrating both pro-inflammatory and resolving responses. We found evidence of local activation of B cells correlated with an age-associated B cell signature, and of progressive stages of monocyte differentiation within the kidney. A clear interferon response was observed in most cells. Two chemokine receptors, CXCR4 and CX3CR1, were broadly expressed, pointing to potential therapeutic targets. Gene expression of immune cells in urine and kidney was highly correlated, suggesting urine may be a surrogate for kidney biopsies. Our results provide a first comprehensive view of the complex network of leukocytes active in lupus nephritis kidneys.

---

Immune-mediated injury of the kidney, referred to as lupus nephritis (LN), is a frequent complication of systemic lupus erythematosus (SLE)^1, 2^. Current immunosuppressive therapies for LN are both toxic and insufficiently effective, and a substantial number of patients progress to end stage renal disease and death^2, 3^. Despite the rapid pace of immunologic discovery, most clinical trials of rationally designed therapies have failed in both general SLE and LN, and only one new drug has been approved for the treatment of SLE in the last 5 decades^2, 4^. Thus, there is a pressing need to decipher the immune mechanisms that drive target organ damage in LN.

To meet this goal, we set out to identify the types and activation states of immune cells in the kidneys of LN patients, as part of the NIH Accelerating Medicines Partnership (AMP) RA/SLE consortium, with the eventual goal of collecting and analyzing immune responses in hundreds of patients. Here, we report the development of a standardized protocol to process kidney samples acquired across a distributed clinical and research network, and to analyze them using single-cell transcriptomics. We applied this pipeline to study a proof-of-concept cohort composed of LN patients and healthy living allograft donors (LD). Our analysis delineates, for the first time, the complex network of leukocytes active in LN kidneys. In addition, we analyzed gene expression in single cells isolated from urine samples of LN patients, showing that these have the potential to be used as surrogates for kidney biopsies in assessing the molecular activation state of renal immune cells.

## RESULTS

### Isolation and processing of LN and LD kidney cells for single-cell transcriptomics

In order to establish a uniform pipeline to analyze kidney biopsy samples acquired at multiple institutions, we evaluated several strategies for their preservation and transport. Cryopreservation of intact, viable kidney tissue offered a method to preserve samples rapidly after acquisition, transport samples with flexible scheduling, and perform sample processing at a central site. We evaluated the feasibility of this method using kidney tissue obtained from tumor nephrectomy samples and then validated the developed workflow using LN and LD biopsies. Kidney tissue samples cryopreserved in a 10% DMSO-containing solution provided robust leukocyte yields, with ∼50% reduction compared to freshly dissociated tissue (Supplementary Fig. 1a). Cryopreserved intact kidney samples yielded higher leukocyte counts than samples that were cryopreserved after tissue dissociation, and comparable leukocyte yields and frequencies to samples shipped overnight in a saline solution on wet ice (Supplementary Fig. 1b,c). Leukocytes from cryopreserved kidney tissue samples showed intact staining for lineage markers by flow cytometry and yielded bulk RNA-seq transcriptomes of equivalent quality to those from non-frozen samples (Supplementary Fig. 1d,e). These results demonstrate that high quality flow cytometric and transcriptomic data can be obtained from leukocytes derived from cryopreserved kidney tissue.

We implemented this pipeline of viable tissue cryopreservation, followed by batched dissociation, cell sorting, and single cell transcriptomics to analyze kidney biopsies from 24 LN patients (Fig. 1a; Supplementary Table 1). A total of 3,541 leukocytes and 1,621 epithelial cells were sorted from LN kidney samples. Ten control samples were acquired from LD kidney biopsies prior to implantation in the recipient, with 438 leukocytes and 572 epithelial cells sorted from these samples. Approximately half of the LN samples provided leukocyte yields well above those obtained from control tissue samples (Supplementary Fig. 2a). High cell yields from LN samples did not segregate with proliferative or membranous glomerulonephritis histologic classification (Supplementary Fig. 2b). Flow cytometric analysis revealed B cells, T cells, macrophages, and other leukocytes in LN samples (Supplementary Fig. 2c,d).

**Figure 1.**
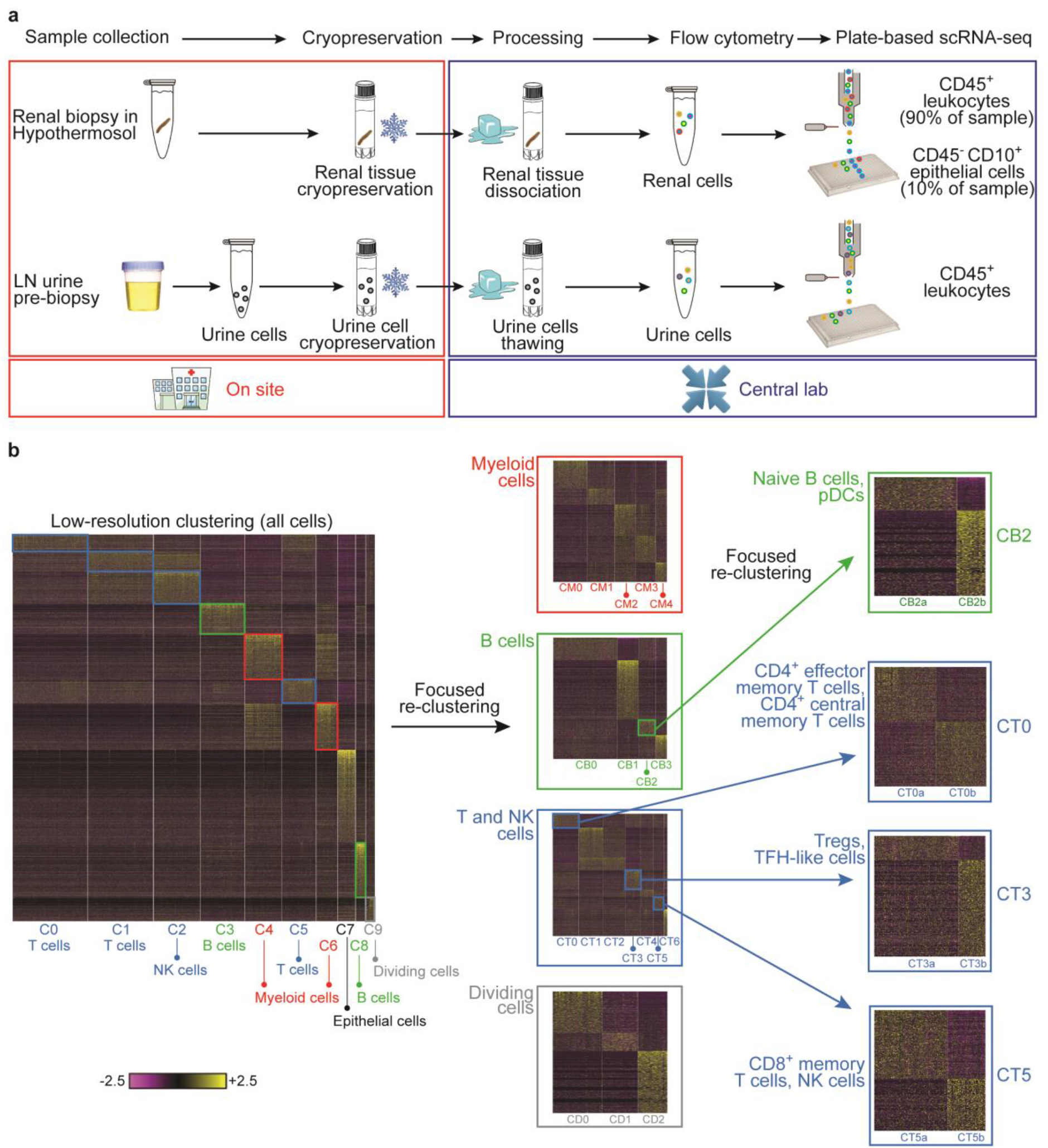
An overview of the approach used for analyzing the cellular contents and molecular states of kidney and urine samples. **a**, Pipeline for collecting and processing kidney and urine samples. Both types of samples were frozen upon collection, then shipped to a central processing cite, in order to minimize batch effects. **b**, Step-wise clustering of kidney cells. Initially, all cells were analyzed together (left heatmap), and the identified clusters were labeled as containing either myeloid cells (red), B cells (green), T/NK cells (blue), dividing cells (gray) or epithelial cells. Each lineage, with the exception of epithelial cells, was then analyzed separately (middle heatmaps), to identify finer clusters. One B cell cluster and three T cell clusters were further re-clustered separately, to generate an even finer description of cell subsets (right).

Viable cells were sorted into 384-well plates, and single-cell RNA-seq (scRNA-seq) was performed using a modified version of the CEL-Seq2 protocol, followed by sequencing of ∼1 million paired-end reads per cell. Since our focus was on characterizing the immune cells active in LN kidneys, 90% of the cells sequenced from each sample were CD45^+^ cells, and the rest were CD45^-^CD10^+^ cells. To exclude low-quality cells and doublets, we filtered out cells that had less than 1,000 or more than 5,000 detected genes, or in which the percentage of mitochondrial genes was larger than 25%. The quality of the collected sequencing data was comparable across plates, and higher in leukocytes compared to epithelial cells, reflecting the lower viability of the latter in the processed samples (Supplementary Fig. 1f-g). Principal component analysis performed on the gene expression data from the remaining 2,736 leukocytes and 145 epithelial cells indicated that the main sources of variability in the data corresponded to cell types, rather than batch or technical factors (Supplementary Fig. 2e-f).

### Step-wise cell clustering identifies cells of the myeloid, T, NK, B and epithelial lineages

In order to identify the lineage of the cells extracted from kidney samples and to characterize their activation state, we clustered them based on their gene expression data, taking a stepwise approach (Fig. 1b). Starting with low-resolution clustering of all kidney cells, we identified 10 clusters (Supplementary Fig. 2g), which we labeled as myeloid cells (clusters C4 and C6), T/NK cells (C0, C1, C2 and C5), B cells (C3, C8), dividing cells (C9) and kidney epithelial cells (C7). Cluster labeling was based on the expression of canonical lineage markers and other genes specifically upregulated in each cluster. Extensive sensitivity analysis demonstrated that while the specific number of clusters at this stage varied with changes in clustering parameters, the assignment of cells to a general lineage or state (myeloid, T/NK, B, epithelial or dividing cells) was highly robust (Supplementary Table 2).

To further resolve cell states, we clustered the cells of each lineage separately based on the most variable genes per lineage, and identified 22 clusters - 21 immune cell clusters and a single epithelial cell cluster (Fig. 2a) - each of which containing cells from multiple patients and plates (Fig. 2b; Supplementary Tables 3-4); this indicates that the identified clusters were largely defined by the types and states of cells rather than patient or batch. Most clusters, with the exception of two (described below), were either absent or present at negligible frequency in LD control samples (Fig. 2b).

**Figure 2.**
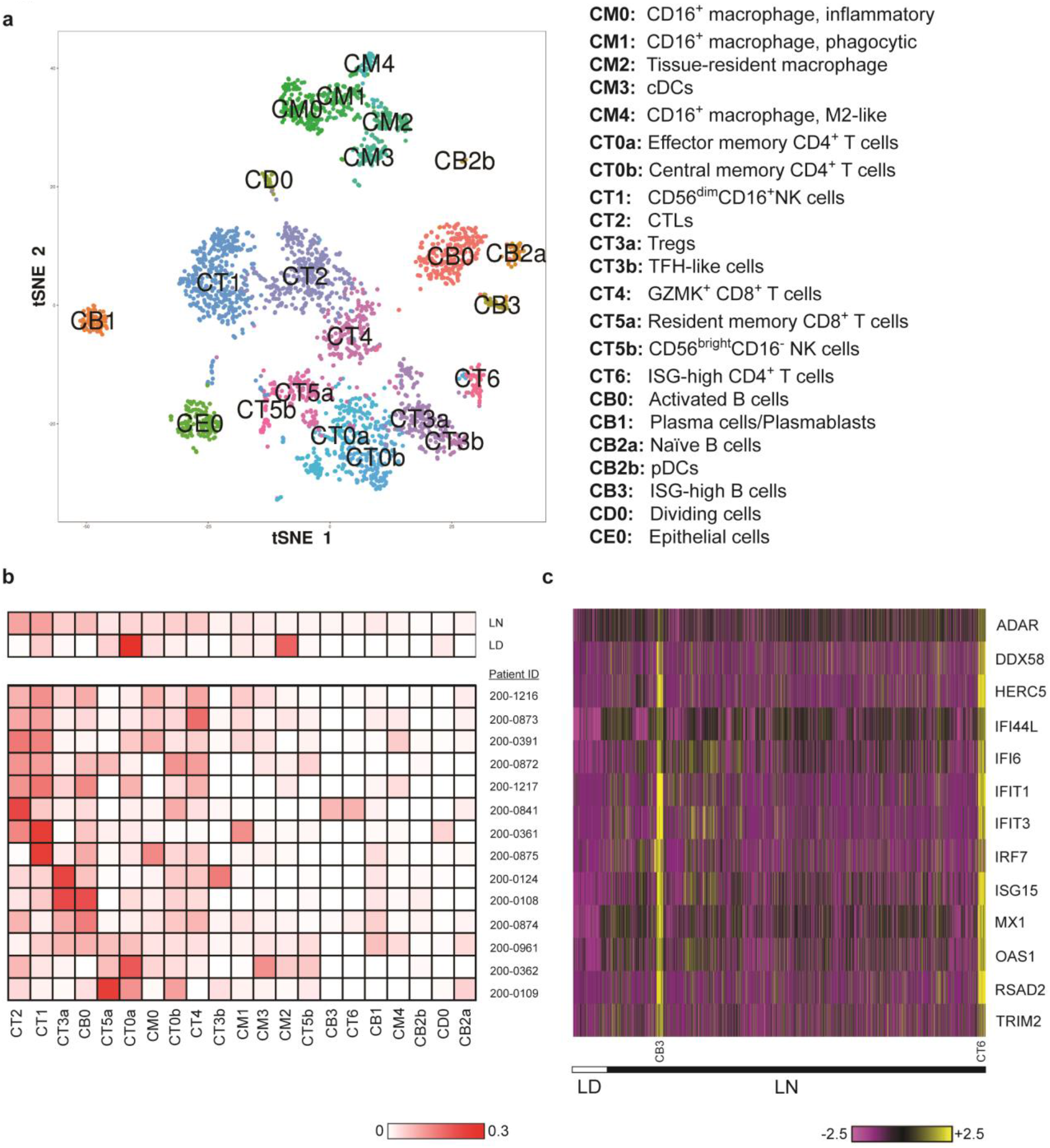
A summary of the step-wise clustering of kidney cells. **a**, 22 clusters were identified; their putative identities are specified on the right. **b**, The fraction of each cluster out of all kidney cells (not including the epithelial cells cluster, CE0), specified for all LN patients and LD controls taken together (top heatmap), and separately for each LN patient that had more than 70 cells passing quality filters (bottom heatmap). **c**, The expression of a selected set of genes induced by type I interferon (ISGs), shown for all kidney cells. The cells of LD controls are displayed on the left side of the heatmap, and all cells are further grouped by cell cluster (separately for LD controls and LN patients). While the expression of ISGs was particularly high in two clusters of cells (CB3 and CT6), these genes were upregulated in almost all cells extracted from LN patients, as compared to LD controls.

Type I interferons have long been known to be elevated in the peripheral blood of lupus patients^5^. In line with this observation, most immune cells in the kidneys of LN patients expressed upregulated levels of interferon-stimulated genes (ISGs), as compared to cells from LD controls (Fig. 2c). Two clusters, one containing B cells and the other CD4^+^ T cells (CB3 and CT6; see below) demonstrated a particularly high expression level of these genes. Interestingly, the majority of cells in these two clusters were extracted from a single patient (patient ID 200-0841), and most of the remaining cells from a second patient (patient ID 200-0874) (Supplementary Table 3). The fact that these two patients also featured B cells and CD4^+^ T cells with a substantially less prominent interferon response suggests that the secretion of this cytokine may be spatially localized to distinct niches, at least in some patients.

### Classification and annotation of myeloid cell clusters reveal resident and infiltrating populations

Focused analysis of the 466 cells in myeloid clusters C4 and C6 revealed 5 finer clusters (clusters CM0-CM4, Fig. 3a; Supplementary Fig. 3a-c). We determined the putative identity of the cells in these clusters by comparing their global gene expression patterns to those of published reference monocyte/dendritic cell (DC) clusters identified in blood samples of healthy individuals using scRNA-seq^6^ (Fig. 3b,c; Supplementary Fig. 3d), and by the expression of canonical lineage markers. Cluster CM3 was closest to CD1c^+^ DCs (reference clusters DC2 and DC3) or Clec9a^+^ DCs (reference cluster DC1), in accordance with the expression of the canonical DC markers CD1C and FLT3, as well as the lack of expression of monocyte markers CD14 and CD16. Cluster CM0 cells were most similar to CD16^+^ patrolling monocytes (reference clusters Mono2 and DC4) with very high expression of CD16 (FCGR3A) and CX3CR1 and low expression of CD14 and CCR2. Clusters CM1 and CM4 were also most similar to reference cluster Mono2; however, CM1 cells expressed lower levels of CX3CR1 and CD16 than CM0, while CM4 cells expressed even lower levels of these two genes and in contrast higher levels of CD14 and CD64 (FCGR1A). These 3 clusters likely represent infiltrating kidney monocyte/macrophage subsets as they constitute a small minority of myeloid cells in normal kidney (Fig. 2b).

**Figure 3.**
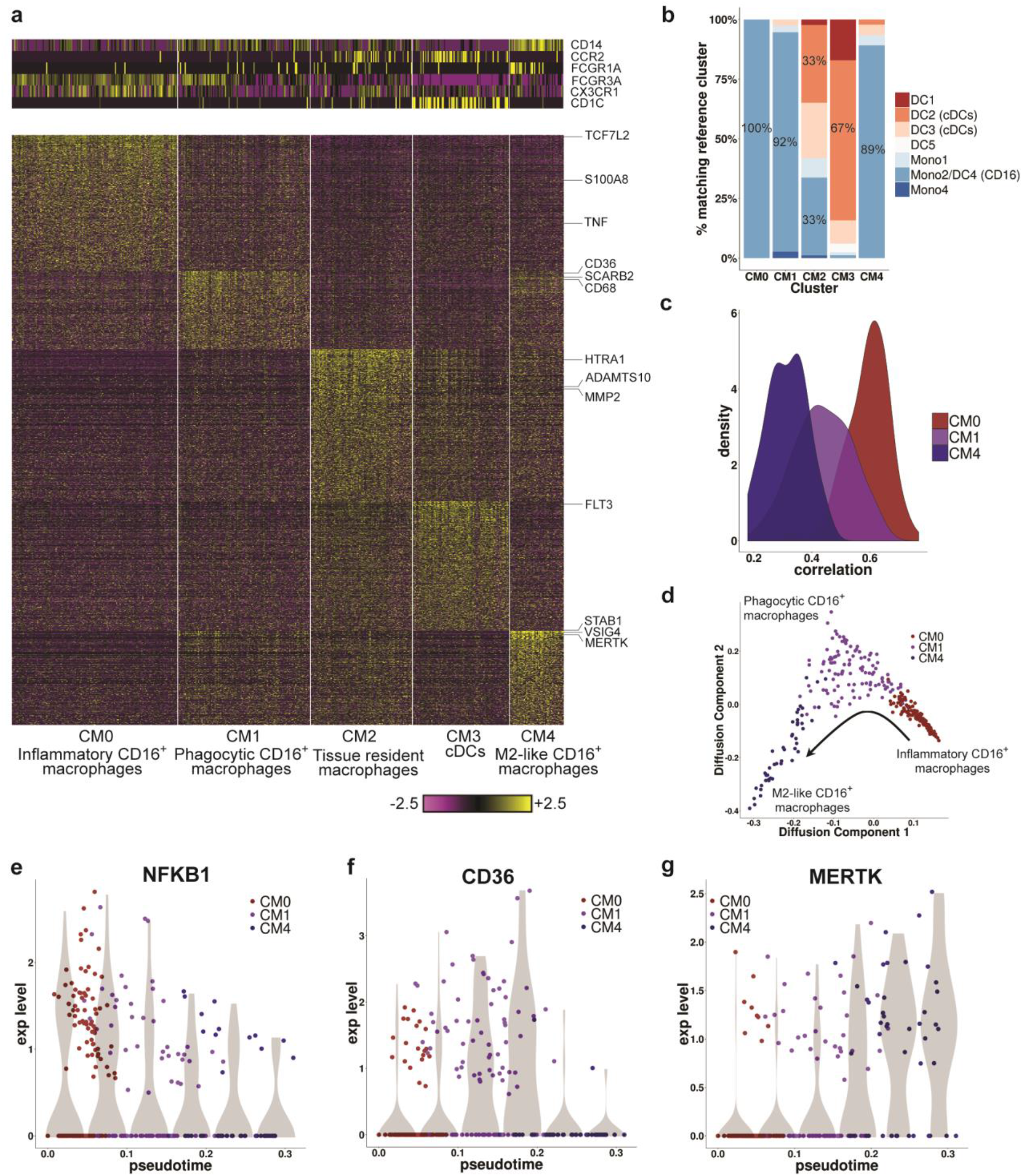
Focused analysis of kidney myeloid cells. **a**, 5 clusters of myeloid cells were identified. The heatmaps show the expression of either canonical lineage markers (top) or genes differentially upergulated in each cluster (bottom). **b**, The results of classifying the kidney myeloid cells by correlating their gene expression to a set of 10 reference clusters (Mono1-Mono4, DC1-DC6), taken from Villani *et al.* For each of the 5 clusters identified in our data, the bars denote the percentage of cells most similar to each of the reference clusters. The percentage of cells mapped to the most frequent reference cluster in each case is specified on the bars. **c**, The distribution of Pearson correlation values, comparing the cells in clusters CM0 (red), CM1 (purple) and CM4 (blue) to the reference clusters DC4 and Mono2 in Villani *et al.* (for each cell, the maximum of the two correlation scores is reported). **d**, The cells of clusters CM0 (red), CM1 (purple) and CM4 (blue), presented in two dimensions using diffusion maps. The arrow represents the direction of the putative transition between these 3 clusters, as explained in the text. **e-g**, The changes in the expression of 3 selected genes, along the trajectory shown in (d); “pseudotime” represents the ordering of the cells along this trajectory. The violin plots (shades) show the distribution of expression levels in equally-spaced intervals along the pseudotime axis (and do not directly correspond to cell clusters).

We next determined whether the pattern of gene expression in each cluster could indicate functional capabilities such as phagocytosis and inflammation, which represent major macrophage activities in damaged tissues^7, 8, 9^ (Supplementary Fig. 3a). Cluster CM1 expressed upregulated levels of phagocytic receptors CD36 (SCARB3), SCARB2, CD68, CD163, NR1H3 (LXR), and GPNMB and cluster CM4 expressed VSIG4, MSR1, CD163, MERTK, STAB1 and CD209. Cells in CM1 and especially CM4 had upregulated expression of C1Q, which not only acts as an opsonin for phagocytosis but also promotes apoptotic cell clearance by enhancing the expression of MERTK (in the presence of other molecules, such as HMGB1) and its soluble ligand GAS6^10, 11^. Clusters CM1 and CM4 also expressed the highest levels of the sialic acid binding immunoglobulin lectin CD169 (SIGLEC1), an endocytic receptor that has been detected on infiltrating macrophages in a variety of human renal inflammatory diseases and that is associated with a phagocytic and reparative phenotype^12^. Cluster CM0 had the highest level of expression of inflammatory genes including TNF, S100A8, S100A9, NFKB1 and the Wnt pathway activator TCF7L2. By contrast, CM4 expressed many genes associated with alternatively activated macrophages, including CD163 and SLC40A1 (ferroportin), which control iron homeostasis^13^. CM4 also abundantly expressed IGF1 and DAB2, both drivers of the alternatively activated phenotype^14, 15^ and folate receptor beta (FOLR2), a receptor expressed on alternatively activated CD14^+^ macrophages that are found in multiple inflammatory and malignant tissues^16^.

Finally, since CM2 was the main cluster found in normal kidneys (Fig. 2b), it likely corresponds to steady-state kidney macrophages. This cluster demonstrated low expression of CD14, CD16, CX3CR1 and CCR2, and no clear similarity to the published reference clusters of peripheral myeloid subsets (Fig. 3b). In comparison to the other macrophage subsets in the kidney, CM2 upregulated several genes associated with tissue remodeling including MMP2, ADAMTS10 and HTRA1. The functions of several other genes preferentially expressed by CM2 are not well defined in macrophages; however, they did express BHLHE41, a gene that is also expressed in microglia and lung resident macrophage populations^17^, consistent with CM2 representing resident cells. Compared to CM2 cells from LD controls, lupus CM2 cells expressed higher levels of interferon stimulated genes, as well as anti-inflammatory genes (GRN, TMSB4X, CREB5) and inhibitors of TLR signaling (GIT2, TNFAIP8L2), and lower levels of proinflammatory genes (ALOX15B, WNT5A) (Supplementary Table 5)^17^.

In summary, a single cluster found in normal and lupus kidneys is likely a resident population (CM2), and four clusters that appeared only in the kidneys of lupus patients - a conventional DCs cluster (CM3) and three CD16^+^ monocyte/macrophage clusters (CM0, CM1, CM4) – are infiltrating myeloid cells.

### Putative transitions between myeloid cell clusters suggest patrolling monocytes become phagocytic or alternatively activated

An important question is how these cell types are related to each other, and whether transitions between the three infiltrating monocyte/macrophage clusters in particular could be inferred from the presence of intermediate states. Indeed, dimensionality reduction using either diffusion maps^18^ (Fig. 3d) or t-Distributed Stochastic Neighbor Embedding^19^ (tSNE; Fig. 2a) indicated possible transitions between these 3 clusters, with CM1 linking CM0 and CM4. Furthermore, since the cells in cluster CM0 tended to be the most similar out of these 3 clusters to peripheral blood CD16^+^ monocytes, while the cells in cluster CM4 were the least similar to their blood counterparts (Fig. 3c), the suggested progression is from an inflammatory blood monocyte (CM0) to a phagocytic (CM1) and then an alternatively activated (CM4) phenotype. As examples of genes reflecting such a progression, we found a gradual reduction along the trajectory from CM0 to CM4 in the expression of NFKB1, an inflammatory gene (Fig. 3e); a transient increase in CD36, an important phagocytic receptor (Fig. 3f); and a continuous increase in MERTK, a key signaling receptor induced by CD36 (Fig. 3g)^20^. We note, however, that other schemes of transitions (or their absence) between these clusters are possible, and further investigation is required to decide between them.

### LN kidneys contain 2 clusters of NK cells and 3 clusters of CD8^+^ T cells

Clusters C0, C1, C2 and C5, comprising 1,764 cells, contained T cells and NK cells. A focused clustering of these cells separated them into 7 finer clusters of NK, CD8^+^ T and CD4^+^ T cells (clusters CT0-CT6, Fig. 4a; Supplementary Fig. 4a). Cluster CT1 contained NK cells, which could be identified by the lack of CD3E and CD3D combined with expression of CD56 (NCAM1) and DAP12 (TYROBP), as well as high expression of cytotoxic genes including PRF1, GZMB, and GNLY. Cluster CT2 featured CD8^+^ T cells expressing a characteristic cytotoxic T lymphocyte (CTL) program including high expression of GZMB, PRF1, and GNLY, similar to the NK cells in cluster CT1. Granzyme B^+^ cells have been identified by immunohistochemistry in LN kidneys and are reported to correlate with more severe disease^21^. A second population of CD8^+^ T cells, demonstrating high levels of a distinct granzyme, GZMK^22^, instead of GZMB and GNLY, populated cluster CT4. These cells expressed relatively low levels of PRF1 compared to cluster CT2 and also showed high expression of HLA-DR/DP/DQ molecules and CCR5, consistent with a prior report^23^. Cluster CT5 contained a mixture of CD8^+^ T cells and NK cells, as indicated by its expression patterns of CD3E, CD3D, CD8A, CD56 and TYROBP. Accordingly, it could be further split into two subclusters (Fig. 4b; Supplementary Fig. 4b): a third CD8^+^ T cell population (cluster CT5a), and a small population of NK cells (CT5b). The cells in cluster CT5a had features of resident memory cells, including expression of ZNF683 (HOBIT), ITGAE, ITGA1, and XCL1 and lack of KLF2^24, 25^. In line with this proposed identity, CT5a was the most abundant CD8^+^ T cell cluster in normal kidney biopsies (Fig. 2b). Cluster CT5b cells expressed TYROBP and CD56, suggesting NK cell identity, and differed from CT1 NK cells by higher expression of KIT, TCF7, IL7R, and RUNX2, and lower expression of PRF1, GZMB, FCGR3A, TBX21, and S1PR5. This expression pattern is consistent with the identification of these cells as tissue-resident CD56^bright^CD16^-^ NK cells, in contrast to the CD56^dim^CD16^+^ NK cell features observed in CT1^26^.

**Figure 4.**
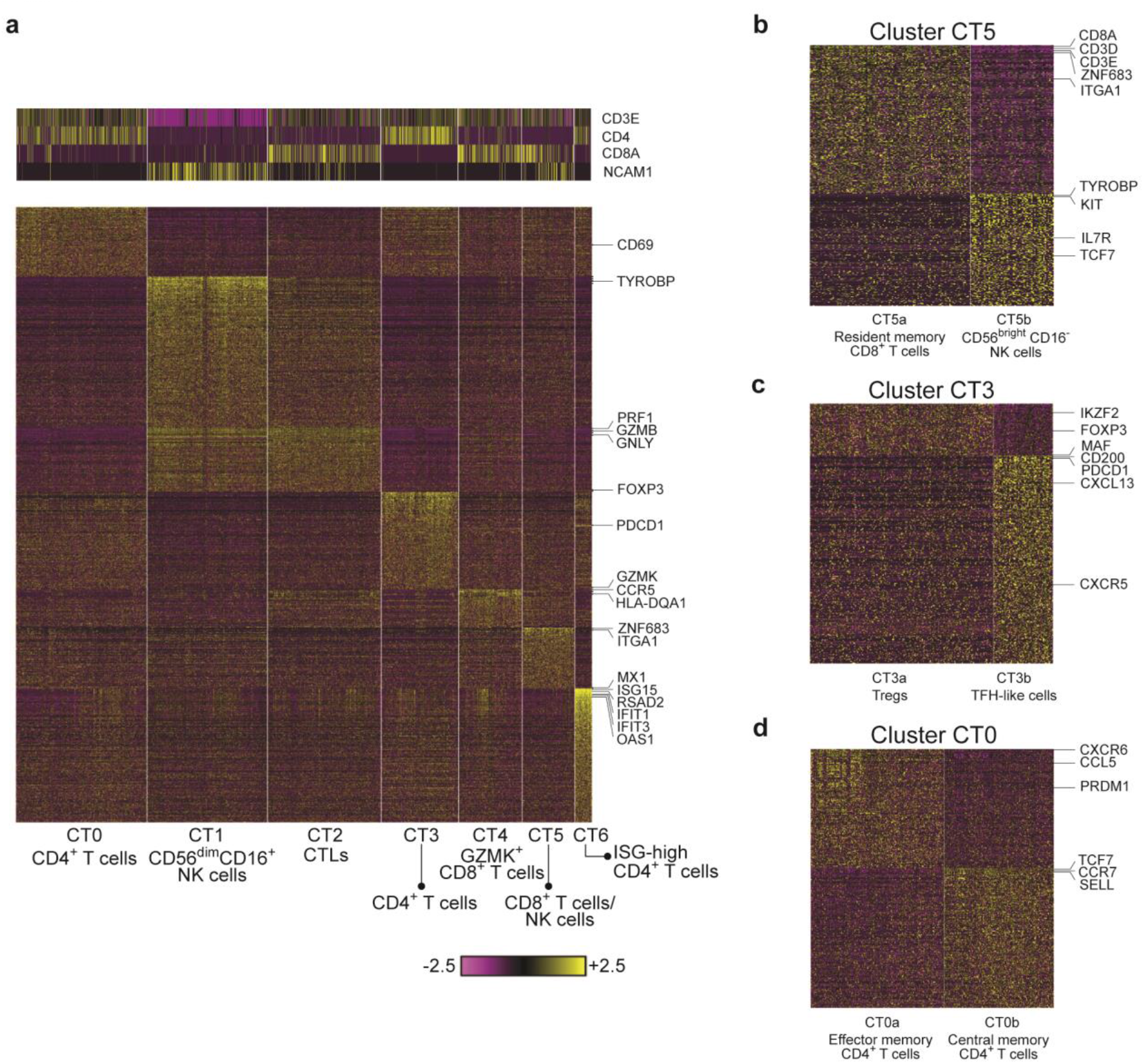
Focused analysis of kidney T cells and NK cells. **a**, Preliminary analysis identified 7 clusters. The heatmaps show the expression of either canonincal markers defining T cell and NK cell subsets (top) or genes differentially upergulated in each cluster specifically (bottom). **b**, The splitting of cluster CT5 into two subclusters, representing resident memory CD8^+^ T cells (CT5a) and CD56^bright^CD16^-^ NK cells (CT5b). **c**, Cluster CT3 can be split into two subclusters, putatively corresopnding to Tregs (CT3a) and TFH-like cells (CT3b). **d**, Subclustering the cells of cluster CT0 reveals two populations of cells, one putatively identified as early effector memory CD4^+^ T cells (CT0a), the other late central memory CD4^+^ T cells (CT0b).

Of note, none of the three CD8^+^ T cell clusters expressed a clear exhaustion signature (Supplementary Fig. 4c), contrary to the identification of such a signature in peripheral blood CD8^+^ T cells in lupus patients, previously reported^27^.

### Analysis of CD4^+^ T cell subsets identifies 5 clusters, including T follicular helper-like cells

Clusters CT0, CT3, CT6 contained CD4^+^ T cells. Cells in cluster CT3 expressed genes associated with T regulatory cells (Tregs), including FOXP3 and IKZF2 (Helios). Interestingly, CT3 also contained cells that expressed genes characteristic of T follicular helper (TFH) cells, including CXCL13, CXCR5, PDCD1, MAF, and CD200 (Fig. 4c; Supplementary Fig. 4d). Indeed, CT3 could be further divided into 2 subclusters, with one subcluster (CT3a) containing CD4^+^ FOXP3^+^ Tregs and a second subcluster (CT3b) consisting of cells with low FOXP3 and features consistent with TFH cells.

Cluster CT0 featured a mixture of CD4^+^ T cells and could be further split into two subclusters, the first containing primarily effector memory CD4^+^ T cells (CT0a), with more frequent expression of PRDM1, CCL5, and CXCR6, and the second consisting of mostly CCR7^+^ SELL^+^ TCF7^+^ central memory T cells (CT0b, Fig 4d; Supplementary Fig. 4e). The similar expression of CD69 in both clusters suggests that CT0b cells are more likely to be central memory than naïve CD4^+^ T cells. Expression of TCF7, KLF2, and LEF1 may indicate an early central memory T cells (Tcm) phenotype of CT0b cells, in contrast to the late effector phenotype of CT0a cells^28^. Interestingly, CT0a was the only CD4^+^ T cell cluster found with substantial frequency in LD samples (Fig. 2b). Comparing gene expression across the two study subject groups, we identified a dysregulated expression of IFN-induced genes in the LN samples (Supplementary Table 5).

Notably, while some LN kidney T cells have been previously annotated as Th1 and Th17 cells, in our data CD4^+^ T cells did not segregate into distinct clusters with characteristic effector lineage features (e.g. Th1, Th2, Th17). IFNG and CXCR3 could be identified in some CT0 cells, primarily within CT0a (Supplementary Fig. 4e). In contrast, IL17A, IL17F, and CCR6 were very rarely detected, and no IL4, IL5, or IL13 expression was observed. TBX21 and RORC were found in a minority of CT0 cells, with TBX21 much more frequently expressed in CD56^dim^CD16^+^ NK cells (cluster CT1) and CTLs (CT2; Supplementary Fig. 4a). Taken together, these results suggest that Th1 cells outnumber Th17 cells in LN kidneys.

Finally, cluster CT6 contained CD4^+^ T cells demonstrating exceptionally higher levels of ISGs, including ISG15, MX1, RSAD2, OAS3, IFIT1 and IFIT2, compared to other T cells (Supplementary Fig. 4a).

### Identification of B cell clusters reveals age-associated B cells

A finer analysis of the 435 cells mapped to clusters C3 and C8 identified 4 different B cell clusters in LN samples, but no B cells in healthy kidneys (clusters CB0-CB3, Fig. 2b, Fig. 5a; Supplementary Fig. 5a). Cluster CB1 clearly contained plasmablasts/plasma cells, expressing high levels of XBP1 and MZB1, as well as immunoglobulin genes. The cells in cluster CB3 demonstrated high levels of several ISGs, including IFIT1, IFIT2, IFIT3, ISG15, OAS3 and RSAD2. Expression of these genes was also detected in the other B cell clusters, but at substantially lower levels.

**Figure 5.**
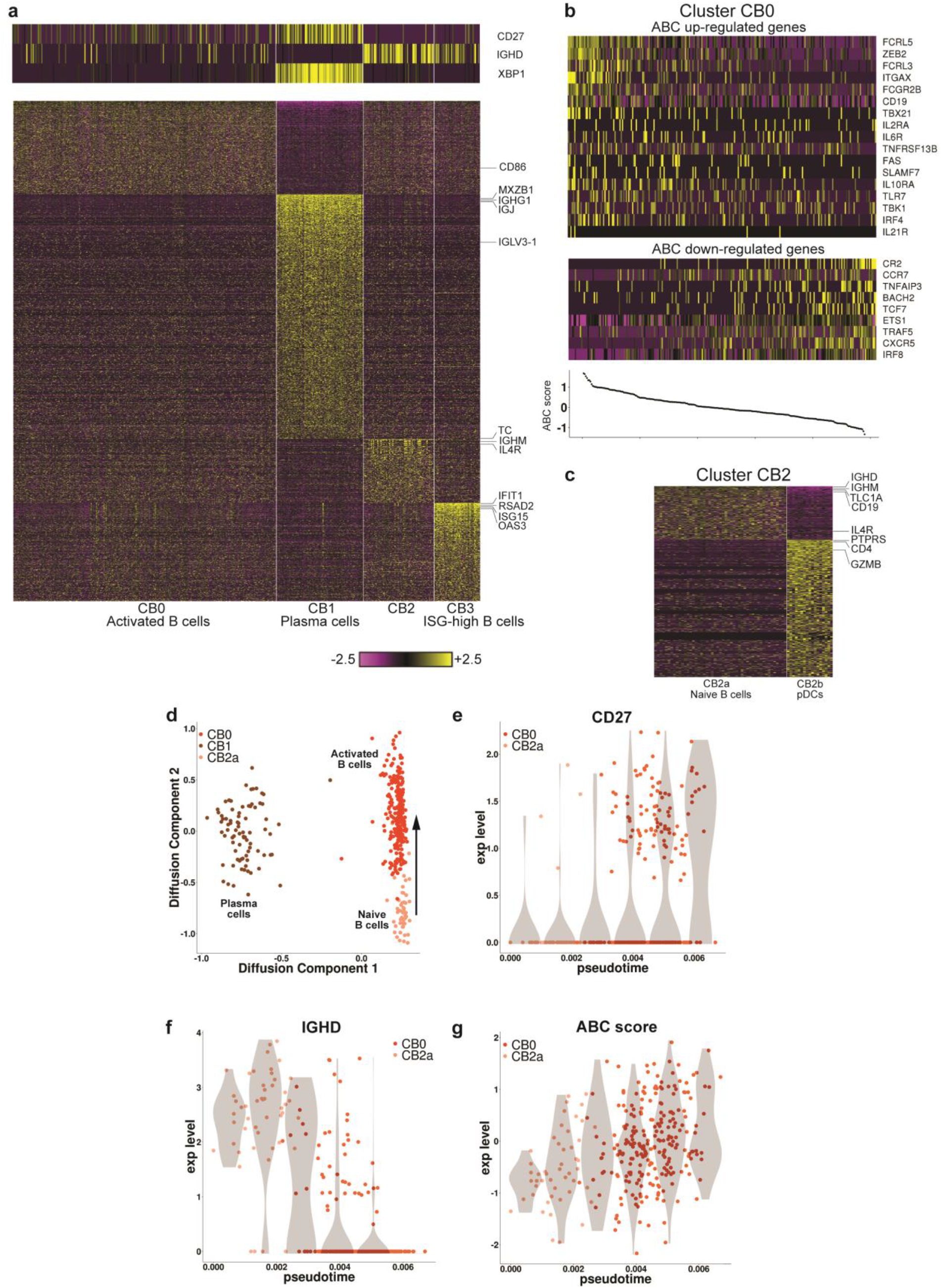
Focused analysis of kidney B cells. **a**, Preliminary analysis identified 4 clusters. The heatmaps show the expression of either canonincal markers defining B cell subsets (top) or genes differentially upergulated in each cluster specifically (bottom). **b**, Cluster CB0 demonostrates a coordinated expression of genes previously found to be differentially expressed in age-associated B cells (ABCs). The top heatmap pertains to genes known to be upregulated in ABCs, the bottom heatmap to genes downregulted in this subset. Columns are sorted by the ABC score, defined as the difference between the average expression of these two sets of genes. A continuous change in this score is observed (bottom plot), implying a continuum of states rather than the presence of a distinct subpopulation of ABCs. **c**, The splitting of cluster CB2 into two subclusters, one corresponding to naïve B cells (CB2a), the other to pDCs (CB2b). **d**, Projection of the cells in clusters CB0, CB1 and CB2a onto a 2-dimensional plane, using diffusion maps. A clear separation of the plasma cells (CB1) from naïve (CB2a) and activated (CB0) cells is observed. In contrast, the latter two populations form a trajectory of state changes. The arrow represents a hypothesized direction of transition along this trajectory, from naïve to activated B cells. **e-f**, The changes in the expression of CD27 and IGHD along the trajectory shown in (d). **g**, The change in the ABC score along this trajectory. In panels e-g, the violin plots (shades) show the distribution of expression levels in equally-spaced intervals along the pseudotime axis (and do not directly correspond to cell clusters).

Cluster CB0 cells had upregulated expression of activation markers such as CD27, CD86, IGJ, IGHG1, and low levels of IGHD and IGHM, observations that point to these cells being activated B cells. Furthermore, we could detect in this cluster a gene expression signature consistent with age-associated B cells (ABCs) (Fig. 5b). These cells are implicated to have a role in both aging and autoimmunity^29^ and express elevated levels of several genes, including CD19, CD11C (ITGAX), TBX21, FCGR2B, IL2RA, IL6R, IL10RA, IL21R, TACI (TNFRSF13B), FAS, SLAMF7, FCRL3, FCRL5, TLR7, TBK1, ZEB2 and IRF4, while downregulating CXCR5, CCR7, CR2, TRAF5, ETS1, TNFAIP3, IRF8, BACH2 and TCF7 (I. Sanz, personal communication). Taking into account both the genes upregulated in ABCs and those downregulated in them, we computed for each cell in cluster CB0 a score representing the extent to which its gene expression pattern matches that expected by an ABC (“ABC score”). A continuous range of values of this score could be observed in cluster CB0, rather than a clear separation into distinct subpopulations of cells (Fig. 5b).

A closer look at the tSNE plot for the B cells suggested that cluster CB2 may contain multiple subsets (Supplementary Fig. 5b). Furthermore, some of the markers distinct to this cluster, such as CD4, PTPRS and GZMB are not known to be expressed in B cells. On further analysis we were able to split the cells in cluster CB2 into two subclusters (Fig. 5c; Supplementary Fig. 5c): the first of these (CB2a), expressing the B cell markers CD19 and CD20 (MS4A1), demonstrated upregulation of genes typical of naïve B cells, including high levels of IGHD, IGHM, TCL1A and IL4R, and had nearly undetectable expression of CD27; the other cluster (CB2b) expressed genes known to be upregulated in pDCs (Fig. 5c; Supplementary Fig. 5c), including PTPRS, GZMB, CLEC4C, CD123 (IL3RA) and CD317 (BST2). To further validate this hypothesized identification, we calculated the Pearson correlation in gene expression between each cell in cluster CB2 and three independent sets of reference samples. These sets included FANTOM5^30, 31^, containing bulk RNA-seq data from 360 cell types, 17 of which are immune cell subsets; bulk RNA-seq data from 13 immune cell populations sorted from healthy individuals (Browne *et al.*, manuscript in preparation); and a scRNA-seq dataset, which includes data from 10 different clusters of dendritic cells and monocytes isolated from healthy blood^6^. This analysis classified all CB2b cells as pDCs, using any of the 3 reference data sets (Supplementary Fig. 5d-f). Furthermore, as predicted almost all of the CB2a cells were classified as naïve B cells, when compared to the data from Browne *et al.* (the only dataset of the three tested that contained multiple B cell populations). These observations lend support to the identification of the CB2a cells as naïve B cells and the CB2b cells as pDCs.

### Indirect evidence for B cell activation and differentiation into age-associated B cells in the kidney

We next asked whether B cell activation and differentiation may take place in the inflamed kidney. Our sequencing data, which is composed of short, 3’ end fragments, is not suitable for a direct analysis of BCR repertoires and their distribution across the B cell clusters. We therefore approached this question looking at indirect evidence, and in particular the existence of intermediate states between the identified B cell subsets. Projecting the gene expression data of the B cells onto 2 dimensions using diffusion maps^18^, we found that the naïve (CB2a) and activated (CB0) B cells formed a continuum of states, demonstrating a gradual increase in CD27 expression and a parallel decrease in IGHD expression, consistent with local activation (Fig. 5d-f). Furthermore, traversing the trajectory from CB2a to CB0 coincided with a continuous increase in the ABC score (Fig. 5g), indicating that activation and differentiation into age-associated B cells are highly correlated processes occurring in LN kidneys. In contrast, very few cells occupied intermediate states between plasma and naïve or plasma and activated B cells, implying that differentiation into plasma cells does not take place in the inflamed kidney. However, as it was previously suggested that age-associated B cells are pre-plasma cells, this question requires a more direct investigation, in particular employing BCR repertoire analysis.

### The dividing cells cluster includes T and NK cells

A focused analysis of cluster C9 found that it can be further partitioned into 3 finer clusters (Supplementary Fig. 6a). The cells in two of these clusters (CD1, CD2) expressed mitochondrial genes and genes associated with a stress response (Supplementary Fig. 6b), indicating lower viability and/or quality. We therefore excluded these cells from subsequent analyses. The cells in cluster CD0 demonstrated upregulated levels of genes participating in cell division, including MKI67, TOP2A, CENPE, CENPF, CDC6, CDC25A, CDK1, SPDL1 and MAPRE1. Classification of these cells by comparison to FANTOM5 indicated the presence of dividing CD8 T cells, NK cells and CD4 Tregs (Supplementary Fig. 6c).

### Cluster-specific expression of genes associated with disease risk

Genome-wide association studies (GWAS) have identified numerous risk alleles and their susceptibility genes in SLE and LN^32, 33^. We analyzed the expression of these genes across the 22 clusters identified in kidneys, and found both expected and surprising cluster-specific expression patterns (Fig. 6a). For example, TLR7, whose role in nucleic acid sensing, B cell activation and differentiation is well established^34, 35^ is expressed in our data in pDCs, myeloid cells and B cells. Lesser known, HIP1, suggested by one report to possibly regulate DC cell immune activity^36^, is expressed in resident memory CD8^+^ T cells (cluster CT5a), CD56^bright^CD16^-^ NK cells (CT5b), cDCs (CM3) and pDCs (CB2b). Another lesser-known gene, LBH, may modulate synovial hyperplasia^37^. Here we found LBH expression in T and B cell subsets, thus raising the possibility that the LBH risk locus impacts both fibroblasts and specific lymphocyte subsets. We also found cluster-specific expression of several poorly annotated SLE susceptibility genes, including WDFY4, CXorf21, and TMEM39A. Interestingly, many of the SLE GWAS genes are transcription factors, supporting the notion that aberrantly regulated gene expression networks in multiple cell types contribute to SLE^38^. Our analysis identified both innate and adaptive immune cell subsets in which these transcription factors (ARID5B, CIITA, ETS, IKZF1, IKZF2, IRF7, IRF8 and PRDM1) are expressed.

**Figure 6.**
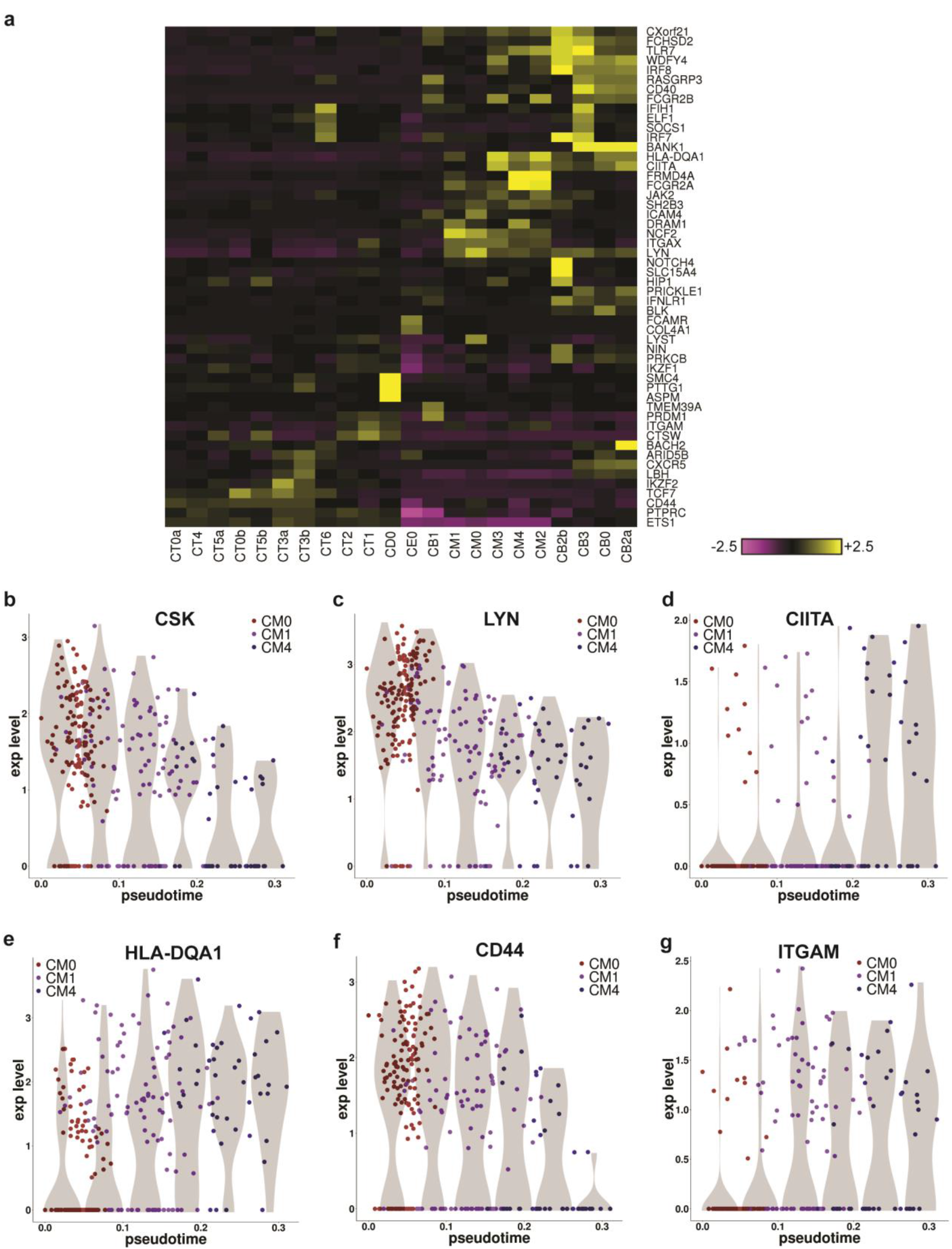
The expression of GWAS genes in lupus nephritis kidneys. **a**, The heatmap shows, for each gene, the scaled average expression across all cells in each cluster. Included are genes previously indicated in lupus by genome-wide association studies, considering only genes that demonstrated high variability across clusters in our data. **b-g**, Changes in the expression of selected GWAS genes, along the trajectory from cluster CM0 to cluster CM4. Violin plots (shades) show the distribution of expression levels in equally-spaced intervals along the pseudotime axis (and do not directly correspond to cell clusters).

We further analyzed the change of some of these genes along the putative trajectory from cluster CM0 to cluster CM4 (Fig. 3d, Fig. 6b-g). We found that CSK and LYN, which regulate the Src family kinase response to weak signals and act as a rheostat for inflammatory signaling^39^ are downregulated along this trajectory, making the macrophages less inflammatory as they transition. A concurrent upregulation of CIITA and HLA-DQA1 suggests an increasing role in antigen presentation. Finally, since it is known that CD44 is required for macrophage infiltration into other organs, the upregulation of ITGAM and downregulation of CD44 raises the hypothesis that these cells become less motile and more adhesive as they transition into the interstitium of the kidney.

### Cellular networks mediated by chemokines and cytokines

To identify signals modulating the activity of leukocytes infiltrating the kidney, we analyzed the expression patterns of chemokine and cytokine receptors. We focused on receptors that were expressed by a relatively large fraction (> 30%) of the cells in at least one cluster (this threshold was set based on the observed distribution of expression frequency, considering all receptors; Supplementary Fig. 7a). Most receptors demonstrated cell type-specific expression patterns (Fig. 7a). However, we found that a single chemokine receptor, CXCR4, was expressed in the majority of infiltrating cells in nearly all clusters (Supplementary Fig. 7b). A second chemokine receptor, CX3CR1, was expressed in most myeloid cells, as well as CD56^dim^CD16^+^ NK cells (cluster CT1) and CTLs (CT2) (Supplementary Fig. 7c). Of note, the expression frequency of other chemokine receptors previously implicated in LN, such as CCR5, CXCR3 and CCR2, was found to be much lower (Supplementary Fig. 7d-f). For cytokine receptors, we observed that IL2RG, encoding the common gamma chain and important for signaling of several cytokines, was frequently expressed in almost all clusters. TGFBR2, a subunit of the receptor for TGFβ was also expressed on the majority of cells. IL10RA, IL27RA, IL17RA and TNFRSF1B were expressed by a large fraction of cells in all clusters, with the exception of the B cells.

**Figure 7.**
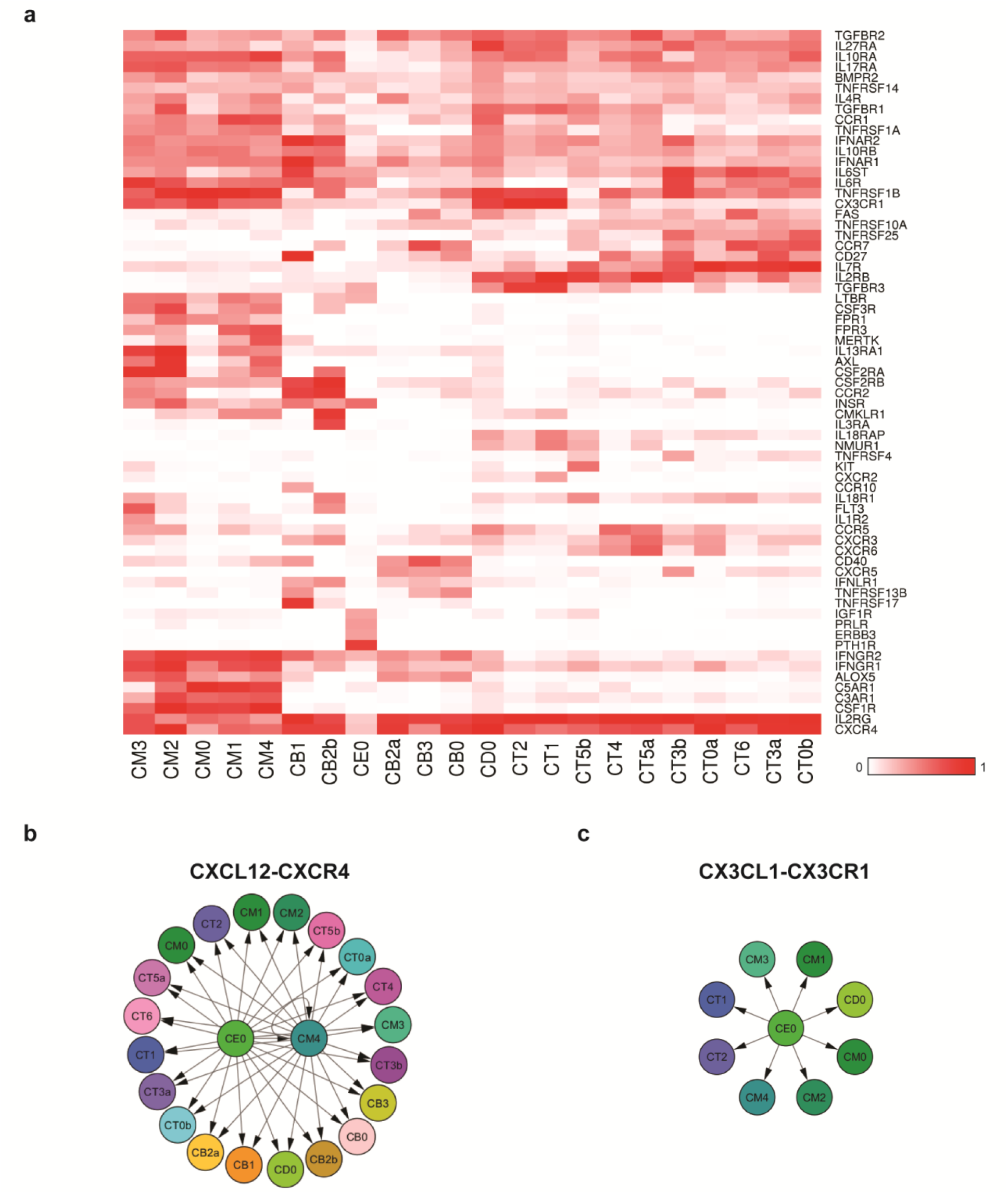
Chemokine- and cytokine-mediated cellular networks. **a**, The pattern of chemokine receptors expression over the cell clusters. The color codes for fraction of cells expressing each receptor. Shown are receptors that are expressed in at least 30% of the cells of at least one cluster. **b**, The producers-consumers cellular network corresponding to the chemokine CXCL12 and its receptor CXCR4. **c**, The producers-consumers cellular network of the chemokine CX3CL1 and its receptor CX3CR1.

By analyzing the expression patterns of the corresponding ligands, one may hope to identify putative interactions between the cells acting in the inflamed kidney. We found that the CXCR4 ligand, CXCL12, was expressed mainly in the cells in cluster CM4, as well as in the epithelial cells (Fig. 7b, Supplementary Fig. 7g). The latter were also the main source of the CX3CR1 ligand, CX3CL1 (Fig. 7c; Supplementary Fig. 7h). Interestingly, CM4 cells were in addition the top producers of CCL2 and CCL8 (Supplementary Fig. 7i), the ligands of CCR2, which is expressed in a high fraction of plasma cells (cluster CB1) and pDCs (cluster CB2b). These findings indicate a possible central role played by the kidney epithelial cells and the M2-like macrophages in cluster CM4 in coordinating the traffic of immune cells infiltrating the kidney.

### Comparison of urine and kidney leukocytes

Leukocytes isolated from urine samples of LN patients were processed in the same way as kidney cells (Fig. 1a). Urine leukocytes tended to have fewer detectable genes, compared to kidney leukocytes (Supplementary Fig. 1h). We therefore applied lower thresholds onto the number of genes detected when discarding low-quality urine cells. Following filtering, 577 high-quality cells, collected from 8 patients, were included in subsequent analyses. Only 3 patients had matching kidney and urine samples with substantial cell numbers; we therefore focused on comparing urine and kidney cells across all patients taken together, rather than performing a patient-based analysis.

We first assigned each urine cell to a cluster of kidney cells, choosing the cluster to which it was most correlated in its gene expression data. We found that compared to kidney, urine samples had an overrepresentation of myeloid cells (in particular cluster CM1) and fewer T cells (Fig. 8a). We next compared gene expression across corresponding urine and kidney clusters, restricting the comparison to clusters that had more than 5 urine cells. High correlations were observed, typically ranging from 0.85 to 0.95 (Fig. 8b). This suggests that gene expression measurements in urine clusters can serve as estimates of kidney gene expression in corresponding clusters; in particular, such estimates were found to be more accurate for genes demonstrating a medium or high expression level (Supplementary Fig. 8a-b).

**Figure 8.**
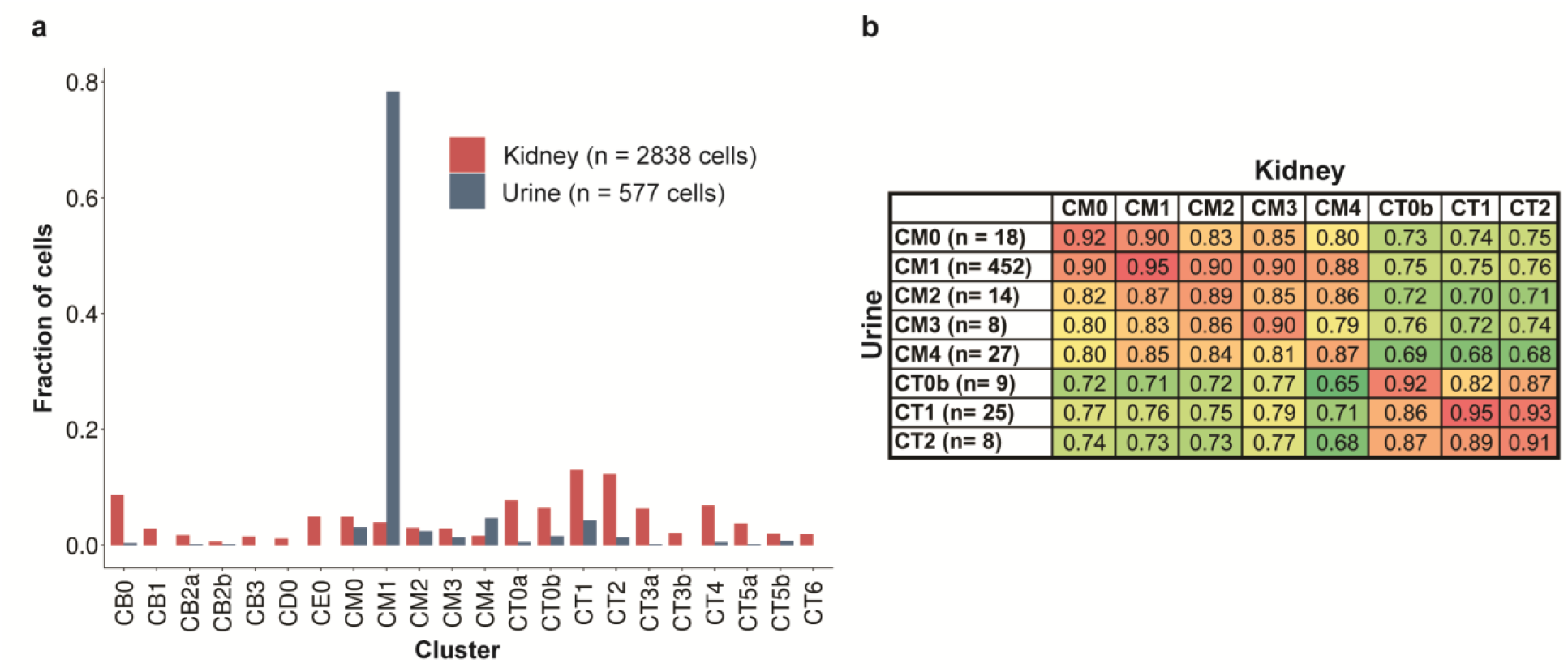
The comparison of cells extracted from urine samples and from kidney samples. **a**, Differences in the relative frequencies of each cluster. **b**, Pearson correlation values between gene expression data of urine and kidney clusters, computed using the average gene expression taken over the cells in each cluster, and considering only clusters that had at least 5 urine cells.

## DISCUSSION

The advent of single cell transcriptomics has enabled analysis of cells from patient samples in unprecedented detail. Here, we used this approach to study, for the first time, kidney samples obtained from LN patients and LD controls. Our analysis revealed the complexity of the immune populations acting in LN kidneys, and showed that these contain several subsets of myeloid, NK, T and B cells, most of which are not found in the kidneys of LD controls. We consider this a first draft map of the immune cell types and states in LN and LD kidneys; sampling of more patients and cells will be required to determine its degree of completeness.

The analysis of T cell clusters revealed several important findings. Remarkably, dividing cells in the kidney were composed almost entirely of T cells and NK cells, suggesting these are actively stimulated and maintained in activated states, consistent with the lack of highly-expressed exhaustion markers in any of the T cell subsets. We identified a large population of CD56^dim^CD16^+^ NK cells that are a major source of IFNγ and of cytolytic molecules (granzyme B, perforin, granulysin); the global transcriptomes will now allow further evaluation of their functional programs. In addition, we identified three distinct populations of CD8^+^ T cells that could not be easily distinguished by surface markers. One population (CT2) appears to be cytotoxic CD8^+^ T cells with expression of effector molecules similar to the NK cells. A second population is distinguished by high expression of granzyme K instead of granzyme B, a pattern that has also been noted in circulating CD8^+^ T cells^22, 23^. Furthermore, we found a resident memory CD8^+^ T cell population, identified based on gene expression similarity to T-RM cells described in other tissues^24, 25^, and present in LD and LN samples. Prior immunohistochemical analyses have revealed CD8^+^ T cells localized to different areas of the kidney^40^; it will therefore be of interest to determine whether the observed CD8^+^ T cell populations, as well as NK cells, show distinct renal localization and functions in autoimmunity.

Our transcriptomic analysis also identified multiple CD4^+^ T cell populations, including a population that expressed features of TFH cells, consistent with prior reports^41, 42^. The clustering together of TFH-like cells and FoxP3^+^ Helios^+^ regulatory T cells further raises the possibility that T follicular regulatory cells are also present^43, 44^. A distinct set of effector CD4^+^ T cells could be subdivided into effector memory and central memory cells. While clusters were not associated with Th1 or Th17 populations, a small number of CD4^+^ T cells did have features of Th1 and rarely Th17 cells, suggesting that T cell polarization may not be a major feature of CD4^+^ T cells in lupus kidneys, and that the bulk of IFNγ is produced by CD8^+^ T cells and NK cells.

B cells are found in more than half of the lupus biopsies but not in healthy samples^45^. The organization of B cell infiltrates can vary from scattered cells, to B-T cell clusters and, rarely, fully formed germinal centers^46, 47^. Clonal expansions of B cells are common, with antigenic specificity frequently directed to local renal antigens^48^, suggesting that immune responses to damaged tissue are being driven *in situ*^49^. Our finding of B cells spanning a spectrum of states between naïve and activated cells, together with the presence of TFH-like cells, is consistent with this view. Interestingly, we found that B cell activation correlates with the expression of molecular features of age-associated B cells (ABCs), a subset of antigen experienced cells that appear to be preferentially driven by BCR/TLR ligands^50^. Understanding whether these cells are clonally expanded or enhance inflammation locally in LN patients, and determining the clonal relatedness of naïve, activated and antibody-secreting B cells will require a larger dataset, combined with analysis of BCR sequences.

Tissue resident myeloid cells have considerable developmental and functional differences from circulating subsets^6, 51^ and their gene expression profile is shaped by their microenvironment^51, 52, 53^. Further changes are induced in cells that phagocytose dead material^54^. In line with this, we identified a putative resident population of renal macrophages that have an interferon signature but also express a number of anti-inflammatory genes.

During tissue injury, monocytes can enter tissues from the peripheral blood and can further differentiate into inflammatory macrophages and reparative/resolving macrophages in situ^55, 56, 57^. If the cells are chronically exposed to damage-associated molecular pattern molecules (DAMPs) and endosomal TLR ligands, resolution may fail, and macrophages with mixed functions may emerge^58^. We identified three distinct sub-populations of infiltrating macrophages, all with a gene expression pattern that most resembled that of peripheral CD16^+^/CD14^dim^ monocytes. These “patrolling” cells function to monitor the integrity of endothelial cells^59^ but can be activated through TLR7 and TLR8 to produce inflammatory cytokines via sensing of nucleic acids produced by damaged cells^60^. Glomerular CD68^+^ macrophages have been reported in many histologic studies of LN, suggesting patrolling behavior, and are associated with more severe clinical disease in a subset of LN patients^60, 61^. Our study suggests that the three subpopulations of CD16^+^ related macrophages identified here may transition through an inflammatory to a resolution phase. Interestingly, CD209 (DC-SIGN), a marker limited to CM4, is known to be expressed predominantly by interstitial macrophages in LN biopsies^62^, showing that this subset has infiltrated the tissue. Expression of CD163, which marks an alternatively activated macrophage phenotype that is highly represented in LN kidneys^63^, was less useful here in distinguishing the macrophage subsets.

DCs may also enter injured kidneys from the peripheral blood, have a short half-life and are most often located within interstitial lymphoid infiltrates^64^. We confirmed that pDCs, myeloid DCs and CLEC9A^+^ DCs are all present in the kidneys of LN patients. Renal infiltration of these classic DC cell types is associated with the later stages of renal injury and fibrosis in human glomerular diseases^64^.

A prominent feature of the gene expression pattern of most of the identified cell types is the expression of IFN inducible genes. While all cells showed this signature to some extent, it was more prominent in some cell types than in others, including subsets of B cells and CD4 T, and varied both between and within patients suggesting that there is spatial heterogeneity with respect to the microenvironment in which this activation occurs.

Our analysis of urine samples showed that while the relative cluster frequencies in the kidney are not accurately reflected in urine, gene expression is highly correlated in these two compartments. This suggests that urine samples can be used to monitor changes over time in the molecular pathways activated in the kidney, in the course of disease and in response to treatment.

Overall, the work presented here demonstrates the feasibility of profiling kidney samples using single cell transcriptomics. Furthermore, our novel strategy to cryopreserve viable kidney tissue immediately after acquisition allowed us to study samples collected at multiple sites, while avoiding major confounding signals due to inter-site or technical variability. A proposed future study, to be performed as part of the AMP consortium, will utilize this strategy to analyze a much larger cohort, in order to investigate how the presence and activation states of particular cell infiltrates are related to disease manifestations and treatment responses. Other extensions will include increasing the number of cells collected from each sample using droplet- or nanowell-based scRNA-seq platforms; sequencing TCRs and BCRs, in order to analyze T and B cell clonality; and profiling stromal renal cells together with leukocytes, in order to elucidate their interactions. The results discovered in such studies will be further validated using tissue staining, functional studies in cell lines or primary human cells, and animal models of disease.

## METHODS

### Human kidney tissue and urine acquisition

Renal tissue and urine specimens from LN patients were acquired at 10 clinical sites in the United States. Institutional review board approval was received at each site. Research biopsy cores were collected from consented subjects either as an additional biopsy pass obtained specifically for research during a clinically-indicated biopsy procedure (9 sites), or as a portion of a biopsy specimen acquired for diagnostic pathology during a clinically-indicated biopsy procedure (1 site). Control kidney samples were obtained at a single site by biopsy of a living donor kidney after removal from the donor and prior to implantation in the recipient.

After acquisition, kidney biopsy samples were placed into HypoThermosol FRS preservation solution for 10-30 minutes on ice and then transferred to a cryovial containing 1 ml of CryoStor CS10 cryopreservation medium (BioLife Solutions). The cryovial was incubated on ice for 20-30 min and was then placed in a Mr. Frosty freezing container (Nalgene, #5100-0001) and transferred to a −80°C freezer overnight. Cryopreserved samples were then stored in liquid nitrogen and shipped on dry ice to the central processing site, where they were stored in liquid nitrogen until processing.

### Kidney tissue thawing and dissociation into a single cell suspension

Kidney samples were thawed and processed in batches of 4 samples, with most batches containing both LN and control kidney samples. The cryovial containing the kidney tissue was rapidly warmed in a 37°C water bath until almost thawed. The sample was then poured into a well of a 24-well dish and rinsed in a second well containing warmed RPMI/10%FBS. The tissue was incubated for 10 minutes at room temperature. Specimens were cut into 2-3 pieces and placed into a 1.5 ml centrifuge tube containing 445uL of Advanced DMEM/F-12 (ThermoFisher Scientific, #12634-028) and 5uL of DNase I (Roche, #04536282001, 100U/ml final concentration). 50uL of Liberase TL (Roche, #05401020001, 250ug/mL final concentration) was added, and the tube was placed on an orbital shaker (300-500 rpm) at 37°C for 12 minutes. At 6 minutes into the digestion, the mixture was gently pipetted up and down several times using a cut 1 mL pipette tip. After 12 minutes, 500uL of RPMI/10% FBS was added to stop the digestion. The resulting cell suspension was filtered through a 70-µm filter into a new 1.7 ml microfuge tube. The cells were washed with RPMI/10%FBS, centrifuged at 300g at 4°C for 10 min, and resuspended in cold PBS for downstream analyses. Quantification of cell yields was performed by hemocytometer with trypan blue exclusion and by flow cytometry with propidium iodide-exclusion. Yields of cell subsets (leukocytes, epithelial cells) were quantified by acquiring the entire sample through the flow sorter and plotting the number of intact, PI-negative cell events with the appropriate surface markers. Cell yields were normalized to input tissue mass.

### Urine cell pellet collection and cryopreservation protocol

Midstream urine samples were collected from LN patients prior to kidney biopsy. The total urine volume (15-90 mL) was split into two 50 mL Falcon tubes. Urine cells were pelleted by centrifugation at 200g for 10 minutes, and then resuspended in 1 ml of cold X-VIVO10 medium (Lonza BE04-743Q). Cells were transferred to a microcentrifuge tube, washed once in 1mL of X-VIVO10 medium, and then resuspended in 0.5 mL of cold CryoStor CS10. Cells were transferred into a 1.8 mL cryovial, placed in a Mr. Frosty freezing container, stored in at −80ºC overnight, and then transferred to liquid nitrogen. For downstream analyses, cryopreserved urine cells were rapidly thawed by vigorous shaking in a 37°C water bath, transferred into warm RPMI/10%FBS, centrifuged at 300g for 10 minutes, and resuspended in cold HBSS/1%BSA.

### Flow cytometric cell sorting

An 11-color flow cytometry panel was developed to identify epithelial cells and leukocyte populations within dissociated kidney cells. Antibodies include anti-CD45-FITC (HI30), anti-CD19-PE (HIB19), anti-CD11c-PerCP/Cy5.5 (Bu15), anti-CD10-BV421 (HI10A), anti-CD14-BV510 (M5E2), anti-CD3-BV605 (UCHT1), anti-CD4-BV650 (RPA-T4), anti-CD8-BV711 (SK1), anti-CD31-AlexaFluor700 (WM59), anti-PD-1-APC (EH12.2H7), and propidium iodide (all from BioLegend). Kidney or urine cells were incubated with antibodies in HBSS/1%BSA for 30 minutes. Cells were washed once in HBSS/1%BSA, centrifuged, and passed through a 70 micron filter. Cells were sorted on a 3-laser BD FACSAria Fusion cell sorter. Intact cells were gated according to FSC-A and SSC-A. Doublets were excluded by serial FSC-H/FSC-W and SSC-H/SSC-W gates. Non-viable cells were excluded based on propidium iodide uptake. Cells were sorted through a 100 micron nozzle at 20 psi. For each sample, 10% of the sample was allocated to sort CD10^+^CD45^-^ epithelial cells as single cells, and the remaining 90% of the sample was used to sort CD45^+^ leukocytes as single cells. Single cells were sorted into 384-well plates containing 0.6µL of 1% NP-40 with index sorting, and plates were immediately frozen and stored at −80 degrees. Flow cytometric quantification of cell populations was performed using FlowJo 10.0.7.

### Library preparation and RNA sequencing

Single cell RNAseq (scRNA-Seq) was performed using the CEL-Seq2 method^65^ with the following modifications. Single cells were sorted into 384-well plates containing 0.6µL 1% NP-40 buffer in each well. Then, 0.6µL dNTPs (10mM each; NEB) and 5nl of barcoded reverse transcription primer (1µg/µL) were added to each well along with 20nL of ERCC spike-in (diluted 1:800,000). Reactions were incubated at 65°C for 5min, and then moved immediately to ice. Reverse transcription reactions were carried out and as previously described^65^ and cDNA was purified using 0.8X volumes of AMPure XP beads (Beckman Coulter). *In vitro* transcription reactions (IVT) were performed as described followed by EXO-SAP treatment. Amplified RNA (aRNA) was fragmented at 80°C for 3min and purified using RNAClean XP beads (Beckman Coulter). The purified aRNA was converted to cDNA using an anchored random primer and Illumina adaptor sequences were added by PCR. The final cDNA library was purified using AMPure XP beads (Beckman Coulter). Paired-end sequencing was performed on the HiSeq 2500 in Rapid Run Mode with a 5% PhiX spike-in using 15 bases for Read1, 6 bases for the Illumina barcode and 36 bases for Read2.

### RNA-seq Data Processing

We used a modified version of the Drop-seq pipeline developed by the McCarroll lab (http://mccarrolllab.com/wp-content/uploads/2016/03/Drop-seqAlignmentCookbookv1.2Jan2016.pdf) adapted for CEL-Seq2, to perform all steps necessary to produce gene by cell expression matrices of reads as well as UMIs. These steps include demultiplexing, quality filtering, polyA and adapter trimming, aligning, and collapsing reads with unique combinations of cell+gene+UMI. We used STAR-2.5.1b to align reads to the Hg19 human genome reference. Only uniquely mapped reads were counted. UMIs with fewer than 10 reads were filtered out before creating the final expression matrices, to minimize reads cross-contamination across cells. For each cell, the computed gene expression counts were then normalized for read depth and log transformed.

### Cell Filtering and Quality Control

High quality cells were defined as having at least 1,000 detected genes (i.e. with positive count values). We further required the percentage of mitochondrial genes per cell to be lower than 25%. To remove wells that were suspected to contain mRNA from multiple cells, we required the number of genes per cell to be smaller than 5,000.

To minimize the effect of technical factors, we tested different regression models, taking into account such variables as the plate ID, number of UMIs per cell, and the percentage of mitochondrial genes per cell. We found that using such models had a negligible effect on the gene by cell expression matrix, as well as the overall results of clustering. We therefore decided to avoid employing them in cleaning the data for subsequent analyses.

### Cell Clustering

Clustering of cells was done using Seurat (v1.4.0.8)^66^, in a step-wise manner. We initially performed low-resolution clustering, analyzing all cells together, then labeled each of the resulting clusters as either myeloid cells, T/NK cells, B cells, dividing cells or epithelial cells. The cells of each such general class were then analyzed separately, in order to identify finer clusters. In some cases, as described in the main text, the resulting clusters further split into subclusters. In each case, clustering was done following PCA, based on context-specific variable genes that were identified independently for each set of analyzed cells.

Sensitivity analysis was performed in each clustering step, with a particular focus on the low-resolution clustering stage. Briefly, all parameters in the clustering process, including the number of variable genes and principal components considered, were varied, and the robustness of the results was determined. To assess this robustness, we computed in each case a consistency score, based on a reference clustering run: looking at a large number (1,000) of random pairs of cells, we counted how many pairs were either included in the same cluster in both of the compared clustering runs, or not included in the same cluster, and referred to these as consistent pairs; we then calculated the fraction of consistent pairs out of all random cell pairs considered. We repeated this procedure 100 times, to calculate the mean consistency score.

### Classification by correlation

In determining their putative identity, we compared the gene expression of individual cells to external gene expression datasets of reference samples. In each comparison, we computed the Pearson correlation between the log-transformed gene expression data of the cell and the reference sample, and chose the reference sample that produced the maximal correlation value.

Myeloid cells were compared to the scRNA-seq data published in Villani *et al*., such that each cluster in that study was represented by the average expression over the cells included in it, taking into account only genes showing high variability in that dataset. Similar results were found if all genes, or only cluster markers, were considered. Comparison to FANTOM5 and the data from Browne *et al.* was done based on the median of reference sample replicates, considering all genes.

### Differential expression analysis

Identification of genes differentially expressed between LN patients and LD controls was done using the framework proposed by McDavid et al.^67^, as implemented by Seurat. P-values were corrected for multiple comparisons using the Benjamini-Hochberg method^68^. DE analysis was performed separately for each cluster with at least 10 cells in both patient groups.

### Trajectory analysis

Trajectory analysis was performed based on dimensionality reduction using diffusion maps, as implemented in the Destiny software package (v2.6.2)^18^. In each case, only the cells relevant to the question at hand were analyzed.

### Calculation of an age-associated B cell score (ABC score)

For each cell in cluster CB0, we defined the ABC score as the average of the scaled (Z-normalized) expression of the genes known to be upregulated in age-associated B cells, minus the average of the scaled expression of the genes known to be downregulated in these cells (see the main text for the lists of upregulated and downregulated genes).

### Analysis of GWAS genes expression

We analyzed the expression patterns of 180 genes previously reported in GWAS studies of either SLE or lupus nephritis. For each such gene, we calculated its average scaled (Z-normalized) expression in each cell cluster, taking into account only cells coming from LN samples. For Bi-clustering of GWAS genes and cell clusters, we kept only genes that had an average scaled expression value of more than 1 or less than −1 in at least one cell cluster, such that bi-clustering was based only on the GWAS genes that were relatively variable in our data. Bi-clustering was then performed, based on the average scaled expression in each cell cluster and using a Euclidean distance metric.

### Analysis of chemokine/cytokine receptors

Analysis of chemokine/cytokine receptors was based on a receptor-ligand pairs list downloaded from the IUPHAR/BPS database (http://www.guidetopharmacology.org/download.jsp) and extended manually to incorporate a number of missing, previously published pairs. For each receptor, we calculated the percentage of cells expressing it in each cell cluster, where a receptor was said to be expressed by a cell if it had at least one mapped read (the results reported here were found to be robust to changes in this threshold). For bi-clustering of receptors and cell clusters, we kept only receptors that appeared in at least 30% of the cells in at least one cluster.

### Assignment of urine cells to kidney clusters

For each urine cell, we computed its Pearson correlation with each kidney cluster, taking the average over the kidney cells included in the cluster. The urine cell was then assigned to the cluster that produced the highest correlation value.

### Code availability

All R scripts used to analyze the data reported in this publication are available from the corresponding authors upon request.

### Data availability

The data reported in this publication are deposited in the ImmPort repository and accessible with accession code SDY997.

## ACKNOWLEDGEMENTS

This work was supported by the Accelerating Medicines Partnership (AMP) in Rheumatoid Arthritis and Lupus Network. AMP is a public-private partnership (AbbVie Inc., Arthritis Foundation, Bristol-Myers Squibb Company, Foundation for the National Institutes of Health, Lupus Foundation of America, Lupus Research Alliance, Merck Sharp & Dohme Corp., National Institute of Allergy and Infectious Diseases, National Institute of Arthritis and Musculoskeletal and Skin Diseases, Pfizer Inc., Rheumatology Research Foundation, Sanofi, and Takeda Pharmaceuticals International, Inc.) created to develop new ways of identifying and validating promising biological targets for diagnostics and drug development. Funding was provided through grants from the National Institutes of Health (UH2-AR067676, UH2-AR067677, UH2-AR067679, UH2-AR067681, UH2-AR067685, UH2-AR067688, UH2-AR067689, UH2-AR067690, UH2-AR067691, UH2-AR067694, and UM2-AR067678). N.H. was supported by the David P. Ryan, MD Endowed Chair in Cancer Research. We thank participating Lupus Nephritis Trials Network clinical sites and participants.

## AUTHORS CONRIBUTIONS

A.A., D.A.R, A.D., P.J.H., A.H.J. and D.J.L. analyzed the data; D.A.R., C.C.B., T.M.E., E.P.B., J.A.L. and D.A.H. developed the sample collection and processing protocols; D.A.R., Y.L., A.C., A.N., D.S. and S.S. processed the samples; S.L., D.J.L., A.N., D.S. and S.S. developed the scRNA-seq library preparation protocol; D.E.S., P.T., E.M., M.D.E., M.P., D.L.K., R.A.F., F.P.S., J.P.B., M.A.P., C.P., K.C.K., E.S.W., D.A.H., D.W. and J.H.A. acquired kidney samples; A.A., D.A.R., C.C.B., A.D., W.A., J.A.L., D.A.H., C.N., D.W., M.K., J.H.A., M.B.B., N.H. and B.D. designed the study; A.D., W.A., D.A.H., C.N., D.W., M.K., J.H.A., M.B.B., N.H. and B.D. supervised the work; A.A., D.A.R., C.C.B., A.D., N.H. and B.D. wrote the manuscript.

## COMPETING INTERESTS

M.B.B. is a consultant to Roche in the area of stromal cells.

